# Lineage-specific rediploidization is a mechanism to explain time-lags between genome duplication and evolutionary diversification

**DOI:** 10.1101/098582

**Authors:** Fiona M. Robertson, Manu Kumar Gundappa, Fabian Grammes, Torgeir R. Hvidsten, Anthony K. Redmond, Sigbjørn Lien, Samuel A.M. Martin, Peter W. H. Holland, Simen R. Sandve, Daniel J. Macqueen

**Affiliations:** Institute of Biological and Environmental Sciences, University of Aberdeen, Aberdeen AB24 2TZ, United Kingdom.; Centre for Integrative Genetics (CIGENE), Faculty of Biosciences, Norwegian University of Life Sciences, Ås NO-1432, Norway.; Department of Chemistry, Biotechnology and Food Science, Norwegian University of Life Sciences, 1432 Ås, Norway.; Umeå Plant Science Centre, Department of Plant Physiology, Umeå Plant Science Centre, Umeå University, SE-90187 Umeå, Sweden.; Centre for Genome-Enabled Biology & Medicine (CGEBM), University of Aberdeen, Aberdeen AB24 2TZ, United Kingdom.; Department of Zoology, University of Oxford, South Parks Road, Oxford, OX1 3PS, United Kingdom.

**Author notes:** **Corresponding author**: Daniel J. Macqueen. **Email addresses:** Fiona M. Robertson, Manu Kumar Gundappa, Fabian Grammes, Torgeir R. Hvidsten, Anthony K. Redmond, Sigbjørn Lien, Samuel A.M. Martin, Peter W. H. Holland, Simen R. Sandve, Daniel J. Macqueen.

**Keywords:** Whole genome duplication, rediploidization, species radiation, <u>L</u>ineage-specific <u>O</u>hnologue <u>Re</u>solution (LORe), duplicate genes, functional divergence, autotetraploidization, salmonid fish

## Abstract

The functional divergence of duplicate genes (ohnologues) retained from whole genome duplication (WGD) is thought to promote evolutionary diversification. However, species radiation and phenotypic diversification is often highly temporally-detached from WGD. Salmonid fish, whose ancestor experienced WGD by autotetraploidization ~95 Ma (i.e. ‘Ss4R’), fit such a ‘time-lag’ model of post-WGD radiation, which occurred alongside a major delay in the rediploidization process. Here we propose a model called ‘Lineage-specific Ohnologue Resolution’ (LORe) to address the phylogenetic and functional consequences of delayed rediploidization. Under LORe, speciation precedes rediploidization, allowing independent ohnologue divergence in sister lineages sharing an ancestral WGD event. Using cross-species sequence capture, phylogenomics and genome-wide analyses of ohnologue expression divergence, we demonstrate the major impact of LORe on salmonid evolution. One quarter of each salmonid genome, harbouring at least 4,500 ohnologues, has evolved under LORe, with rediploidization and functional divergence occurring on multiple independent occasions > 50 Myr post-WGD. We demonstrate the existence and regulatory divergence of many LORe ohnologues with functions in lineage-specific physiological adaptations that promoted salmonid species radiation. We show that LORe ohnologues are enriched for different functions than ‘older’ ohnologues that began diverging in the salmonid ancestor. LORe has unappreciated significance as a nested component of post-WGD divergence that impacts the functional properties of genes, whilst providing ohnologues available solely for lineage-specific adaptation. Under LORe, which is predicted following many WGD events, the functional outcomes of WGD need not appear ‘explosively’, but can arise gradually over tens of Myr, promoting lineage-specific diversification regimes under prevailing ecological pressures.

## Background

Whole genome duplication (WGD) has occurred repeatedly during the evolution of vertebrates, plants, fungi and other eukaryotes (reviewed in [1-4]). The prevailing view is that despite arising at high frequency, WGD is rarely maintained over macroevolutionary (i.e. Myr) timescales, but that nonetheless, ancient WGD events are over-represented in several species-rich lineages, pointing to a key role in long-term evolutionary success [1, 5]. WGD events provide an important source of duplicate genes (ohnologues) with the potential to diverge in protein functions and regulation during evolution [6-7]. In contrast to the duplication of a single or small number of genes, WGD events are unique in allowing the balanced divergence of whole networks of ohnologues. This is thought to promote molecular and phenotypic complexity through the biased retention and diversification of highly interactive signalling pathways, particularly those regulating development [8-10].

As WGD events dramatically reshape opportunities for genomic and functional evolution, it is not surprising that an extensive body of literature has sought to identify causal associations between WGD and key episodes of evolutionary history, for example species radiations. Such arguments are appealing and have been constructed for WGD events ancestral to vertebrates [11-15], teleost fishes [16-19] and angiosperms (flowering plants) [10, 20-22]. Nonetheless, it is now apparent that the evolutionary role of WGD is complex, often lineage-dependent and without a fixed set of rules. For example, some ancient lineages that experienced WGD events never underwent radiations, including horseshoe crabs [23] and paddlefish [e.g. 24], while other clades radiated explosively immediately post-WGD, for example the ciliate *Paramecium* species complex [25]. In addition, apparent robust associations between WGD and the rapid evolution of species or phenotypic-level complexity may disappear when extinct lineages are considered, as documented for WGDs in the stem of vertebrate and teleost evolution [26-27].

Such findings either imply that the causative link between WGD and species radiations is weak, or demand alternative explanations. In the latter respect, it is has become evident that post-WGD species radiations commonly arise following extensive time-lags. For example, major species radiations occurred >200 Myr after a WGD in the teleost ancestor (‘Ts3R’) ~320-350 Ma [3, 28-29]. In angiosperms, similar findings have been reported in multiple clades [30-31]. Such findings led to the proposal of a ‘WGD Radiation Lag-Time’ model, where some, but not all lineages within a group sharing ancestral WGD diversified millions of years post-WGD, due to an interaction between a functional product of WGD (e.g. a novel trait) and lineage-specific ecological factors [30]. Within vertebrates, salmonids provide a text-book case of a delayed species radiation post-WGD, where a role for ecological factors has been strongly implied [32]. In this respect, salmonid diversification was strongly associated with climatic cooling and the evolution of a life-history strategy called anadromy [32] that required physiological adaptations (e.g. in osmoregulation [33]) enabling migration between fresh and seawater. Importantly, a convincing role for WGD in such cases of delayed post-WGD radiation is yet to be demonstrated, weakening hypothesized links between WGD and evolutionary success. Critically missing in the hypothesized link between WGD and species radiations is a plausible mechanism that constrains the functional outcomes of WGD from arising for millions or tens of millions of years after the original duplication event. Here we provide such a mechanism, and uncover its impact on adaptation.

Following all WGD events, the evolution of new molecular functions with the potential to influence long-term diversification processes depends on the physical divergence of ohnologue sequences. This is fundamentally governed by the meiotic pairing outcomes of duplicated chromosomes during the cytogenetic phase of post-WGD rediploidization [11, 34-35]. Depending on the initial mechanism of WGD, rediploidization can be resolved rapidly or be protracted in time. For example, after WGD by allotetraploidization, as recently described in the frog *Xenopus leavis* [36], WGD follows a hybridization of two species and recovers sexual incompatibility [11]. The outcome is two ‘sub-genomes’ within one nucleus that segregate into bivalents during meiosis [35]. In other words, rediploidization is resolved instantly, leaving ohnologues within the sub-genomes free to diverge as independent units at the onset of WGD. The other major mechanism of WGD, autotetraploidization, involves a spontaneous doubling of exactly the same genome. In this case, four identical chromosome sets will initially pair randomly during meiosis, leading to genetic exchanges (i.e. recombination) that prohibit the evolution of divergent ohnologues and enable an ongoing ‘tetrasomic’ inheritance of four alleles [35]. Crucially, rediploidization may occur gradually over tens of millions of years after autotetraploidization [35, 37].

Salmonid fish provide a vertebrate paradigm for delayed rediploidization post-autotetraploidization (reviewed in [37]). The recent sequencing of the Atlantic salmon *(Salmo salar* L.) genome revealed that rediploidization was massively delayed for one-quarter of the duplicated genome and associated with major genomic reorganizations such as chromosome fusions, fissions, deletions or inversions [38]. In addition, there are still large regions of salmonid genomes that behave in a tetraploid manner in extant species [e.g. 38-40], despite the passage of ~95 Myr since the Ss4R WGD [32]. In light of our understanding of salmonid phylogeny [32, 41], we can also be certain that rediploidization has been ongoing throughout salmonid evolution [38] and was thus likely occurring in parallel to lineage-specific radiations [32, 42]. However, the outcomes of delayed rediploidization on genomic and functional evolution remain uncharacterized in both salmonids and other taxa. In the context of the common time-lag between WGD events and species radiation, this represents a major knowledge gap. Specifically, as explained above, a delay in the rediploidization process will cause a delay in ohnologue functional divergence, theoretically allowing important functional consequences of WGD to be realized long after the original duplication.

Here we propose ‘Lineage-specific Ohnologue Resolution’ or ‘LORe’ as a mechanism to address the role of delayed rediploidization on the evolution of sister lineages sharing an ancestral WGD event (Fig. 1). It builds on and unifies ideas/data presented by Macqueen and Johnston [32], Martin and Holland [43] and Lien et al. [38] and is a logical outcome when rediploidization and speciation events occur in parallel. Under LORe, the rediploidization process is not completed until after a speciation event, which will result in the independent divergence of ohnologues in sister lineages (Fig. 1). This leads to unique predictions compared to a ‘null’ model, where ohnologues started diverging in the ancestor to the same sister lineages due to ancestral rediploidization. Consequently, under LORe, the evolutionary mechanisms allowing functional divergence of gene duplicates [6-7, 11] become activated independently under lineage-specific selective pressures (Fig. 1). Conversely, under the null model, ohnologues share ancestral selection pressures, which hypothetically increases the chance that similar gene functions will be preserved in different lineages by selection (Fig. 1). A phylogenetic implication of LORe is a lack of 1:1 orthology when comparing ohnologue pairs from different lineages (Fig. 1), leading to the past definition of the term ‘tetralog’ to describe a 2:2 homology relationship between ohnologues in sister lineages [43]. Thus, LORe may be mistaken for small-scale duplication if the underlying mechanisms are not appreciated. Despite this, LORe ohnologues have unique phylogenetic properties (Fig. S1) and can be distinguished from small-scale gene duplication by their location within duplicated (or ‘homeologous’) blocks on distinct chromosomes sharing collinearity [38, 44-45].

**Fig. 1.**
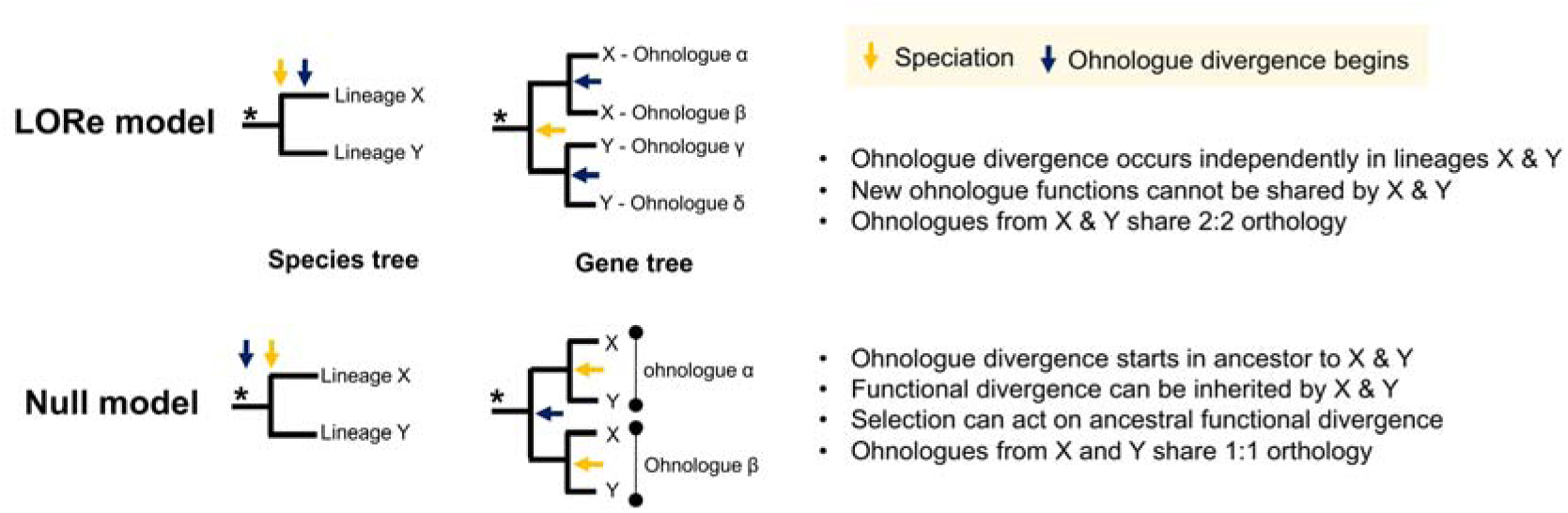
The LORe model of post-WGD evolution following delayed rediploidization. This figure describes the phylogenetic predictions of LORe in contrast to the null model, as well as associated connotations for functional divergence and sequence homology relationships.

In this study, we demonstrate that LORe has had a major impact on the evolution of salmonid fishes at multiple levels of genomic and functional organization. Our findings allow us to propose that LORe, as a product of delayed rediploidization, offers a general mechanism to explain time-lags between WGD events and subsequent lineage-specific diversification regimes.

## Results

### Extensive LORe followed the Ss4R WGD

To understand the extent and dynamics of lineage-specific rediploidization in salmonids, we used in-solution sequence capture [46] to generate a genome-wide ohnologue dataset spanning the salmonid phylogeny [32, 41]. Note, here we use the term ohnologue, but elsewhere ‘homeologue’ has been used to describe gene duplicates retained from the Ss4R WGD event [38]. In total, 384 gene trees were analysed, sampling the Atlantic salmon genome at regular intervals, and including ohnologues from at least seven species spanning all the major salmonid lineages, plus a sister species (northern pike, *Esox lucius)* that did not undergo Ss4R WGD [47] (Table S1). All the gene trees included verified Atlantic salmon ohnologues based on their location within duplicated (homeologous) blocks sharing common rediploidization histories [38]. Salmonids are split into three subfamilies, Salmoninae (salmon, trout, charr, taimen/huchen and lenok spp.), Thymallinae (grayling spp.) and Coregoninae (whitefish spp.), which diverged rapidly ~45 and 55 Ma (Fig. 2). Hence, phylogenetic signals of LORe are evidenced by subfamily-specific ohnologue clades (Fig. 1; Fig. S1). In accordance with this, our analysis revealed a consistent phylogenetic signal shared by large duplicated blocks of the genome, with 97% of trees fitting predictions of either the LORe or the null model (Fig. 3; Table S1; Text S2). The LORe regions represent around one-quarter of the genome and overlap extensively with chromosome arms known to have undergone delayed rediploidization [38].

**Fig. 2.**
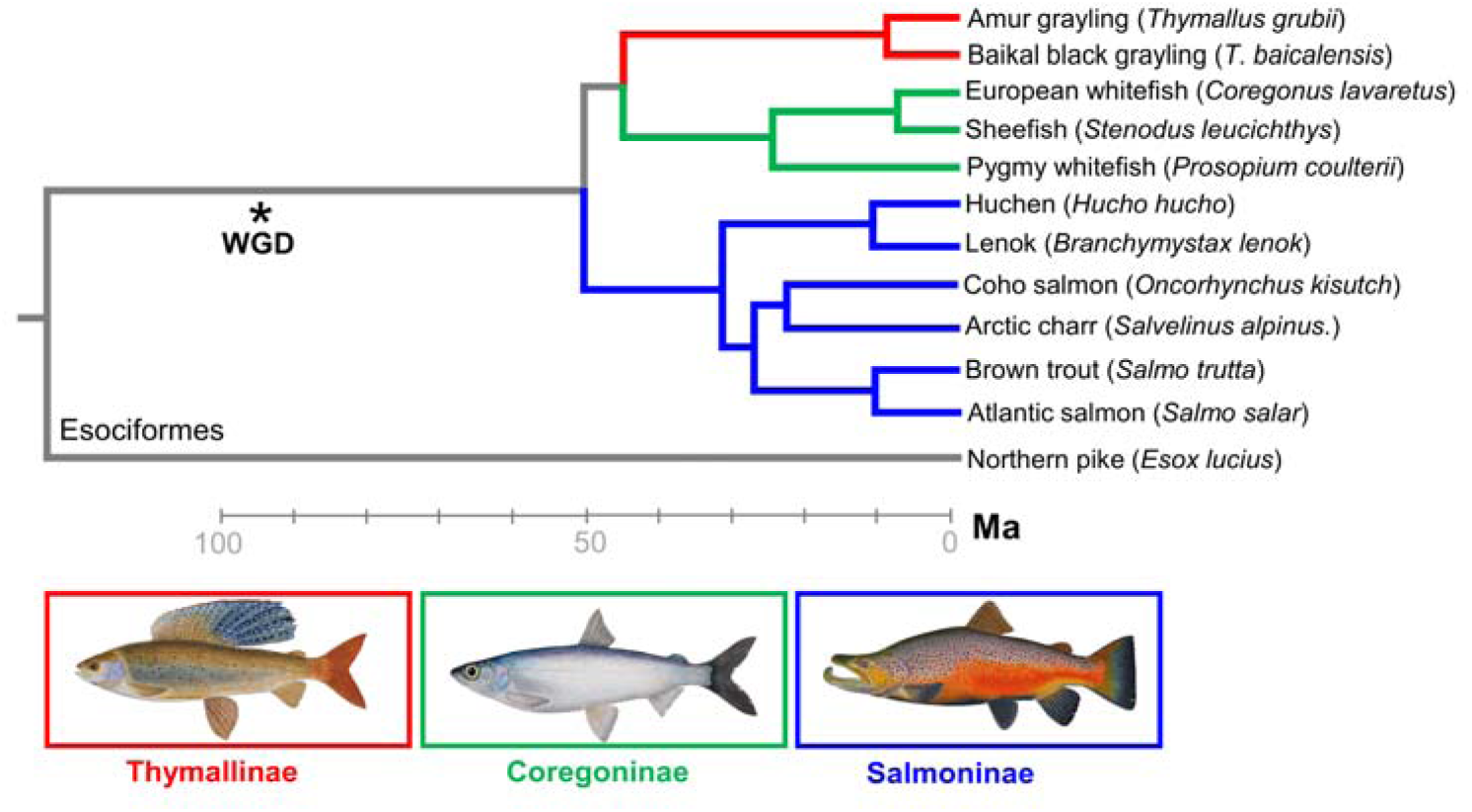
Time-calibrated salmonid phylogeny (after [32]) including the major lineages used for sequence capture and phylogenomic analyses of ohnologues.

**Fig. 3.**
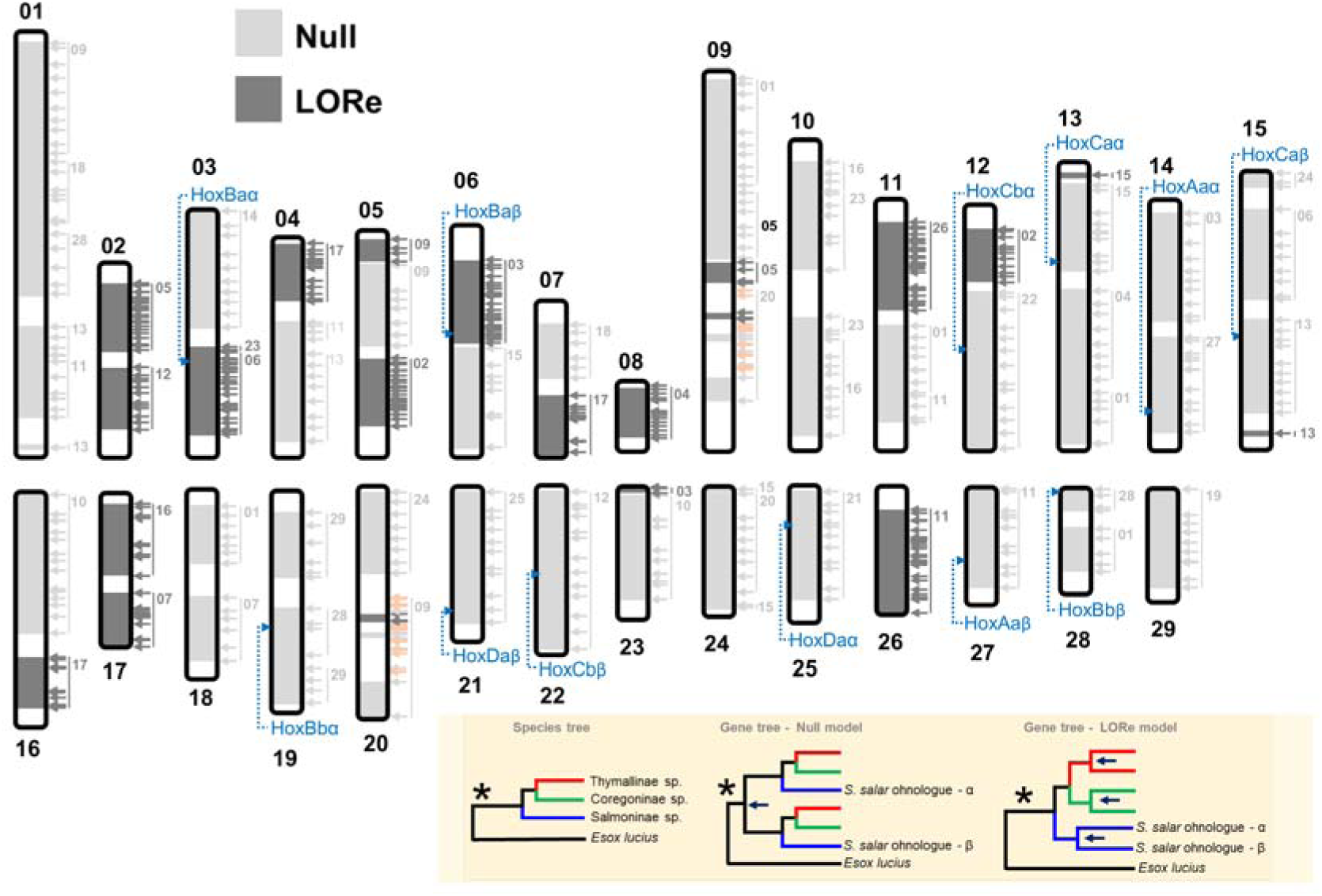
Genome-wide validation of LORe in salmonids. Atlantic salmon chromosomes with LORe and null regions of the genome are highlighted, based on sampling 384 separate ohnologue trees (data in Table S1). Each arrow shows a sampled ohnologue tree (light grey: null; dark grey: LORe; orange: ambiguous; see Text S2). The other chromosome in a pair of collinear duplicated blocks [38] is highlighted, along with the genomic location of salmonid Hox clusters. The shaded box shows the phylogenetic topologies used to draw conclusions about the LORe vs. null model in contrast to other scenarios (Fig. S1).

To complement this genome-wide overview (Fig. 3), we performed a finer-resolution analysis of Hox genes included in our sequence capture study. Hox genes are organized into genomic clusters located across multiple chromosomes and have been used to confirm separate WGD events in the stem of the vertebrate, teleost and salmonid lineages [43, 48-49]. Hox clusters (HoxBa) residing within predicted LORe regions in Atlantic salmon (Fig. 3) showed a highly congruent LORe signal, both considering individual gene trees and trees built from combining separate ohnologue alignments sampled within duplicated Hox clusters (e.g. Fig. 4A; Text S1; Figs S2-S10). Our data indicates that two salmonid-specific Hox cluster pairs underwent rediploidization as single units, either once independently in the common ancestor of each salmonid subfamily for HoxBa (Fig. 4A) or twice in Coregoninae for HoxAb (Fig, S9; Text S1). These results cannot be explained by small-scale gene duplication events under any plausible scenario (Text S1). The HoxAa, HoxBb, HoxCa, HoxCb and HoxDa cluster pairs conformed to the null model (e.g. Fig. 2B; Fig. S10; Text S1), as predicted by their genomic location (Fig. 3). Thus, HoxAb and HoxBa clusters were in regions of the genome that remained ‘tetraploid’ until after the major salmonid lineages diverged ~50 Ma (Fig. 2).

**Fig. 4.**
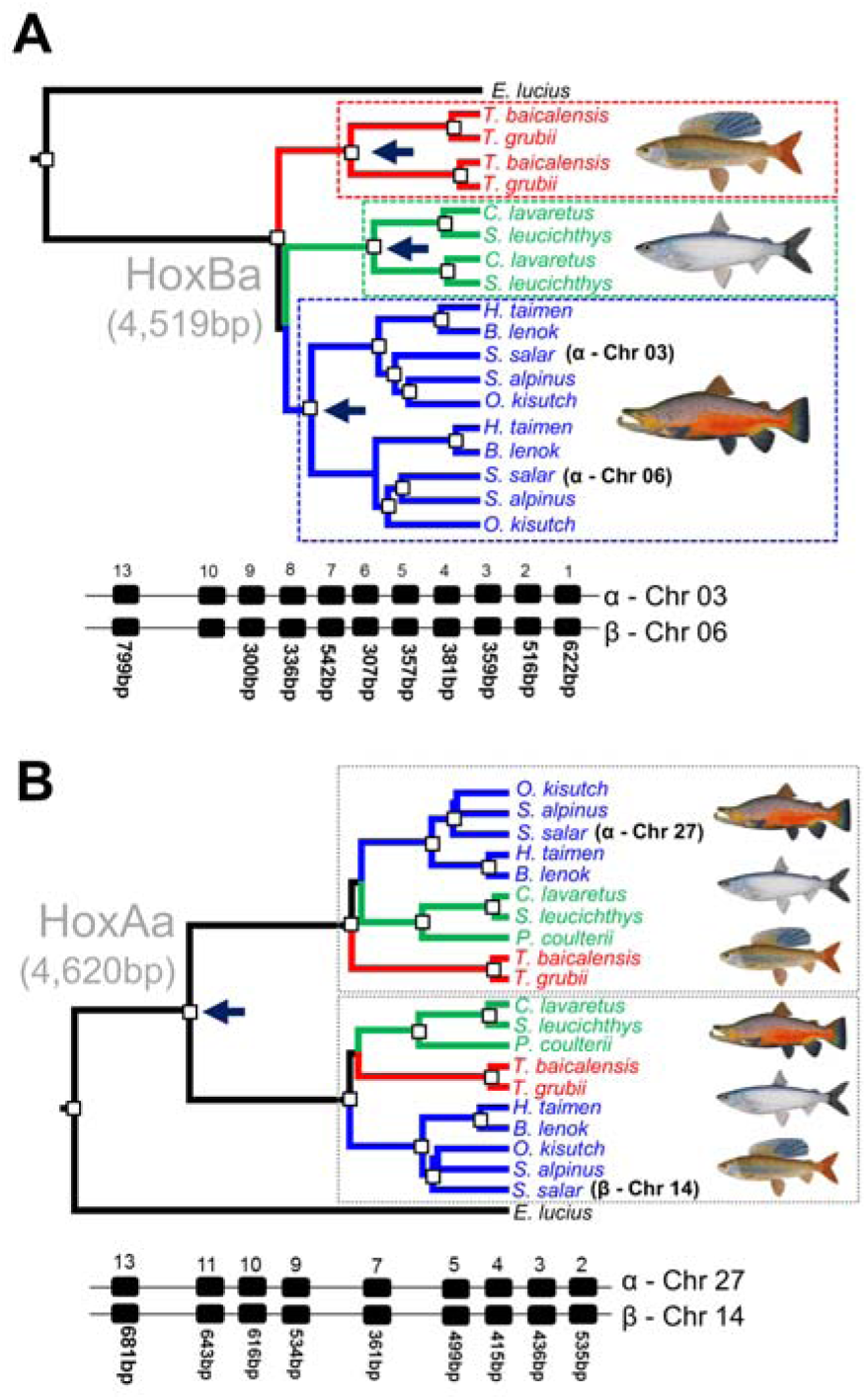
Bayesian phylogenetic analyses of salmonid Hox gene clusters fitting to the predictions of the LORe (**A**) and null (**B**) models. White boxes depict posterior probability values >0.95. Hox clusters characterized from Atlantic salmon [49] are shown, along with the length of individual sequence alignments combined for analysis. The individual gene trees for Hox alignments are shown in the Supplemental Material (Fig. S2/S4 for HoxAa/HoxBa, respectively). Dark blue arrows highlight the inferred onset of ohnologue divergence, i.e. the node where rediploidization was resolved.

We also studied proteins encoded within Hox clusters to contrast patterns of sequence divergence under the LORe and null models (Fig. S12, S13). The data supports our predictions (Fig. 1), as LORe has allowed many amino acid replacements to become independently fixed among Hox ohnologues within each salmonid subfamily (Fig. S12). These changes are typically highly conserved across species, suggesting lineage-specific purifying selection within a subfamily (Fig. S12). Conversely, under the null model, numerous amino acid replacements that distinguish Hox ohnologues arose in the common salmonid ancestor and have been conserved across all the major salmonid lineages (Fig. S13).

### Distinct rediploidization dynamics across salmonid lineages

Our data highlights distinct temporal dynamics of rediploidization across salmonid lineages. For example, using a Bayesian approach, the onset of divergence for the HoxBa-α and -β clusters of Salmoninae, Coregoninae and Thymallinae (i.e. Fig. 4A tree) was estimated at ~46, 25 and 34 Ma, respectively (95% posterior density intervals: 31-57, 15-37 and 21-47 Ma, respectively). Thus, the genomic regions containing these duplicated Hox clusters experienced rediploidization much earlier in Salmoninae than the grayling or whitefish lineages. This is consistent with a broader pattern observed by genome-wide gene tree sampling (Table S1, Fig. 3), which allowed rediploidization events to be mapped along the salmonid phylogeny (Fig. 5A). Based on this data, we estimate that the respective rate of rediploidization in Salmoninae, Thymallinae and Coregoninae was ~59, 39 and 14 ohnologue pairs per Myr, for regions of the genome covered in our analysis. Hence, our data reveal that a relatively higher fraction of LORe ohnologues began to diverge during the early evolution of Salmoninae, leading up to point when anadromy evolved [42], while a much smaller fraction of ohnologues had experienced rediploidization at the same point of whitefish evolution (Fig. 5A). For graylings, the split of the species used in our study represents the crown of extant species, dated at no earlier than 15 Ma [50]. Thus, about two-thirds of LORe ohnologues started diverging in the ancestor to extant graylings (Fig. 5A).

**Fig. 5.**
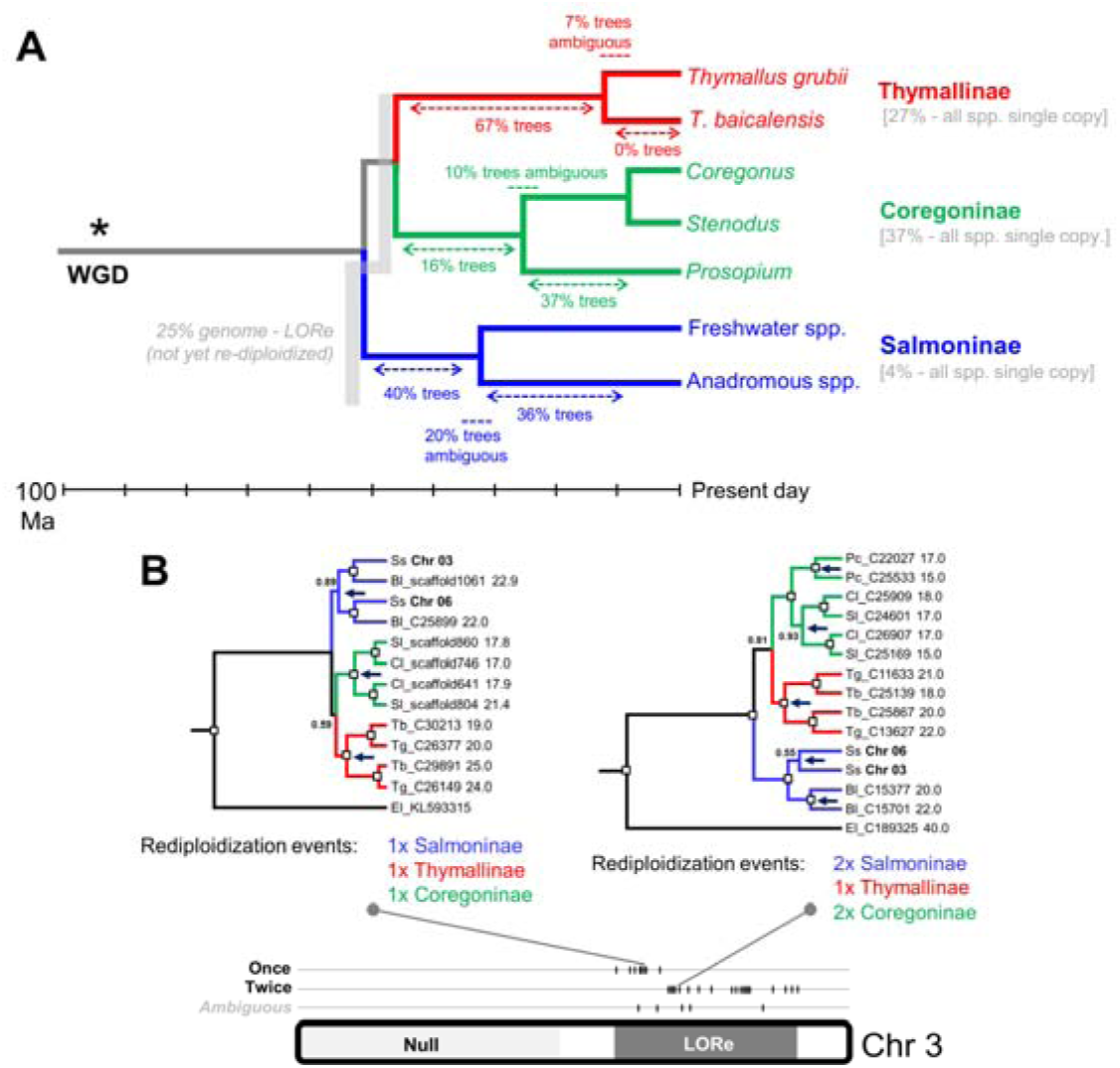
Divergent rediploidization dynamics in different salmonid lineages. (**A**) Time-tree of species relationships [32] showing the fraction of 384 gene trees supporting independent rediploidization events at different nodes (**B**). A LORe region on Chr. 03 (paired with an ohnologous region on Chr. 06), where the number of independent rediploidization events inferred within Salmoninae (shown) is consistent along contiguous regions of the genome. Example trees are shown for genomic regions with distinct rediploidization histories. Abbreviations: Ss: *S. salar*; Bl: *Brachymystax lenok*; Sl: *Stenodus leucichthys;* Cl: *Coregonus lavaretus;* Pc: *Prosopium coulteri*; Tb: *Thymallus baicalensis*; Tg: *T. grubii*.

Interestingly, around one third of our gene trees included a single ohnologue copy for all whitefish and grayling species, which were clustered along chromosomes in the genome (Table S1). As these regions have experienced delayed rediploidization, this likely reflects the ‘collapse’ of highly-similar sequences in the assembly process into single contigs [38], rather than the evolutionary loss of an ohnologue. For two LORe regions with evidence of multiple rediploidization events within a salmonid subfamily, we mapped our findings back to Atlantic salmon chromosomes (Fig. 5B). This showed that the number of inferred rediploidization events within a LORe region is consistent across large genomic regions (Fig. 5B; Fig. S11). Overall, these data support past observations that the rediploidization process is dependent on chromosomal location [38], while emphasizing highly-distinct dynamics of rediploidization in different salmonid subfamilies.

### Regulatory divergence under LORe

To understand the functional implications of LORe, we contrasted the level of expression divergence between Atlantic salmon ohnologue pairs from null and LORe regions (Fig. 6). This was done in multiple tissues under controlled conditions (Fig. 5A, B) and also following ‘smoltification’ [33], a physiological remodelling that accompanies the life-history transition from freshwater to saltwater in anadromous salmonid lineages (Fig. 6C). The regions of the genome covered by our phylogenomic analyses (Fig. 3) contained 16,786 high-confidence ohnologues, with 27.1% and 72.9% present within LORe and null regions, respectively. Ohnologue expression was more correlated within LORe than null regions, both across tissues (Fig. 6B, C) and when considering differences in regulation between fresh and saltwater (Fig. 6C).

**Fig. 6.**
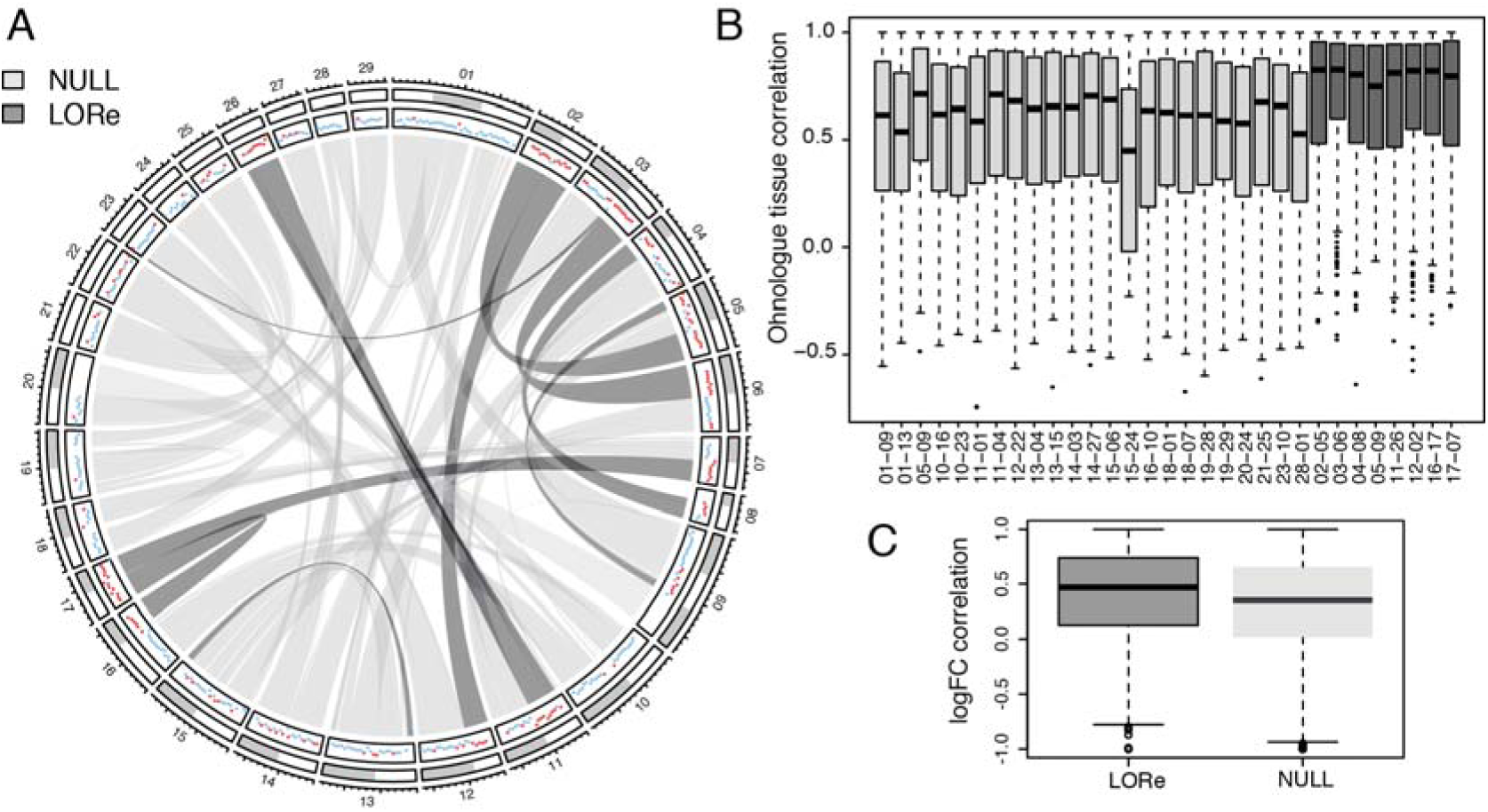
Global consequences of LORe for the evolution of ohnologue expression. (**A**) Circos plot of Atlantic salmon chromosomes highlighting LORe and null regions defined by phylogenomics. The panel with coloured dots indicates expression similarity among ohnologue pairs: each dot represents the correlation of ohnologue expression across a 4 Mb window. Red and blue dots show correlations ≥ 0.6 and <0.6, respectively. (**B**) Correlation in expression levels across 15 tissues for ohnologue pairs in null and LORe regions. Different collinear blocks are shown [38] containing at least ten ohnologue pairs. (**C**) Boxplots showing the overall correlation in the expression responses of ohnologues from LORe and null regions (2,505 and 6,853 pairs, respectively) during the physiological transition from fresh to saltwater. The correlation was calculated for log fold-change responses across 9 tissues.

A recent analysis [38] suggested that 28% of salmonid ohnologues fit a model of expression divergence where one duplicate maintained the ancestral tissue expression (as observed in northern pike) and the other acquired a new expression pattern (i.e. ‘regulatory neofunctionalization’ [38]). We extended this analyses by partitioning ohnologue pairs from LORe and null regions of the genome. Among 2,021 ohnologue pairs displaying regulatory neofunctionalization, ~19% vs. ~81% were located in LORe and null regions, respectively, constituting a significant enrichment in null regions compared to the background expectation (i.e. 27.1% vs. 72.9%) (Hypergeometric test, *P* = 2e-13). The average higher correlation in expression and lesser extent of regulatory neofunctionalization for ohnologues in LORe regions is expected, as they have had less evolutionary time to diverge in terms of sequences controlling mRNA-level regulation. Nonetheless, many ohnologues in LORe regions have clearly diverged in expression (Fig. 6), which may have contributed to phenotypic variation available solely for lineage-specific adaptation.

### Role of LORe in lineage-specific evolutionary adaptation

To better understand the role of LORe in salmonid adaptation, we performed an in-depth analysis of Atlantic salmon genes with established or predicted functions in smoltification [33], hypothesized to represent important factors for the lineage-specific evolution of anadromy. Interestingly, LORe regions contain ohnologues for many genes from master hormonal systems regulating smoltification, including the insulin-like growth factor (IGF), growth hormone (GH), thyroid hormone (TH) and cortisol pathways (Table S2) [33, 51-53]. We also identified LORe ohnologues for a large set of genes involved in osmoregulation and cellular ionic homeostasis, key for saltwater tolerance, including Na+, K+ - ATPases (targets for the above mentioned hormone systems [33, 51]), along with members of the ATP-binding cassette transporter, solute carrier and carbonic anhydrase families (Table S2). Several additional genes from the same systems were represented by ohnologues in null regions (Table S2).

To characterize the regulatory evolution of ohnologues with regulatory roles in smoltification, we compared equivalent tissue expression ‘atlases’ from Atlantic salmon in fresh and saltwater (Fig. 7; Dataset S1). The extent of regulatory divergence was variable for ohnologues in both LORe and null regions, ranging from conserved to unrelated tissue responses (Fig. 7A; Dataset S1). Several pairs of ohnologues from both LORe and null regions showed marked expression divergence in tissues of established importance for smoltification (examples in Fig. 7B, Dataset S1). For example, a pair of LORe ohnologues encoding IGF1 located on Chr. 07 and 17 (i.e. homeologous arms 7q–17qb under Atlantic salmon nomenclature [38]), despite differing by only a single conservative amino acid replacement, were differentially regulated in several tissues (Fig. 5B). The differential regulation of IGF1 ohnologues in gill and kidney is especially notable, as both tissues are vital for salt transport and, in gill, IGF1 stimulates the development of chloride cells and the upregulation of Na+, K+-ATPases, together required for hypo-osmoregulatory tolerance [54-55]. Thus, key expression sites for IGF1 are evidently fulfilled by different LORe ohnologues and these divergent roles must have evolved specifically within the Salmoninae lineage, 40-50 Myr post WGD [32]. In contrast to IGF1, LORe ohnologues for GH, another key stimulator of hypo-osmoregulatory tolerance [33], showed highly conserved regulation during smoltification (Dataset S1). Overall, these findings demonstrate that many Atlantic salmon ohnologues in both LORe and null regions are differentially regulated under a physiological context that recaptures key lineage-specific adaptations for the evolution of anadromy.

**Fig. 7.**
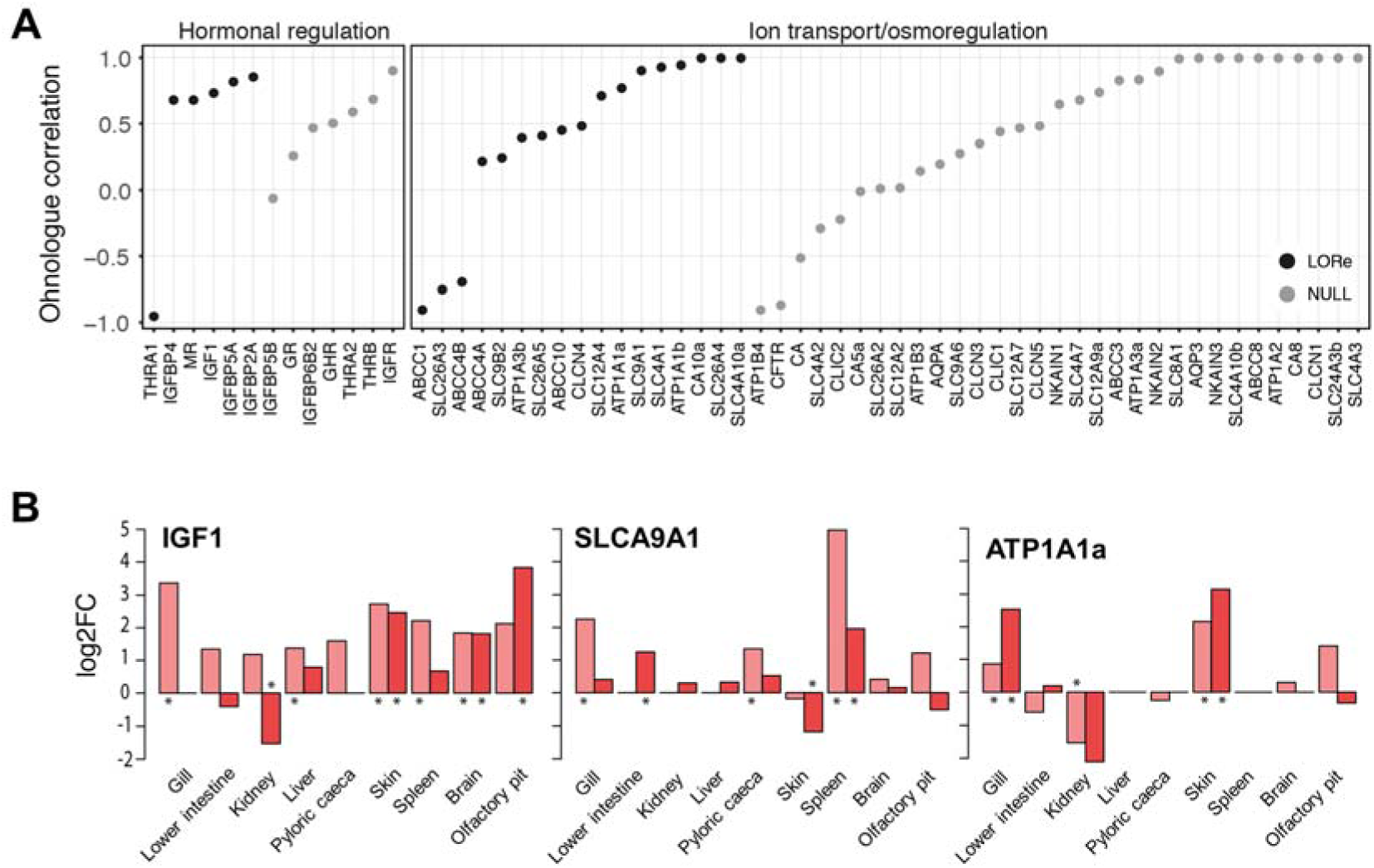
Regulatory evolution of salmonid ohnologues defined within a lineage-specific context of physiological adaptation. (**A**) Correlation in expression responses for ohnologues from LORe vs. null regions during the fresh to saltwater transition in Atlantic salmon. Each name on the x-axis is a pair of ohnologues (details in Table S2). The data is ordered from the most to least correlated ohnologue expression responses. Correlation was performed using Pearson’s method. Data for additional ohnologues where correlation was impossible due to a restriction of expression to a limited set of tissues is provided in Dataset S1. (**B**) Example ohnologues showing a multi-tissue differential expression response to the fresh to saltwater transition. The asterisks highlight significant expression responses. Equivalent plots for all genes shown in part A is provided in Dataset S1.

To further characterize the role of LORe in lineage-specific adaptation, we performed gene ontology (GO) enrichment analysis contrasting all ohnologues present in LORe vs. null regions (Table S3). Remarkably, ohnologues in LORe vs. null regions were enriched for 99.9% non-overlapping GO terms, suggesting global biases in encoded functions (Table S3). The most significantly enriched GO-terms for LORe ohnologues were ‘indolalkylamine biosynthesis’ and ‘indolalkylamine metabolism’ (Table S3). This is notable as 5-hydroxytryptamine is an indolalkylamine and the precursor to serotonin, which plays an important role controlling the master pituitary hormones that govern smoltification [51, 56]. An interesting feature of rediploidization is the possibility that functionally-related genes residing in close genomic proximity (e.g. due to past tandem duplication) started diverging into distinct ohnologues as single units, for example Hox clusters (Fig. 4). We found that the LORe ohnologues contributing to enriched GO-terms ranged from being highly-clustered in the genome, to not at all clustered (Table S4). In the latter case, we can exclude any biases linked to regional rediploidization history. In the former case, we noted that two clusters of globin ohnologues on Chr. 03 and 06 (i.e. homeologous arms 3q-6p under Atlantic salmon nomenclature [38]) explain the enriched term ‘oxygen transport’ (Table S4). This is interesting within a context of lineage-specific adaptation, as haemoglobin subtypes are regulated during smoltification to increase oxygen-carrying capacity and meet the higher aerobic demands of the oceanic migratory phase of the life-cycle [33]. Other GO terms enriched for LORe ohnologues included pathways regulating growth and protein synthesis, immunity, muscle development, proteasome assembly and the regulation of oxidative stress and cellular organization (Table S3).

## Discussion

Here we define the LORe model and characterized its impacts on multiple levels of organization, adding a novel layer of complexity to our understanding of evolution after WGD. While past analyses have highlighted the quantitative extent of delayed rediploidization for a single salmonid genome [38], our study is the first to establish the genome-wide functional impacts of LORe and is unique in revealing divergent rediploidization dynamics across the major salmonid lineages. Our results show that salmonid ohnologues can have strikingly distinct evolutionary ‘ages’, both for different genes located within the same genome (Fig. 3, 4) and when comparing the same genes in phylogenetic sister lineages sharing the same ancestral WGD (Fig. 5). Our data also indicates that thousands of LORe ohnologues have diverged in regulation or gained novel expression patterns tens of Myr after WGD, likely contributing to lineage-specific phenotypes (Fig. 7). Hence, in the presence of highly delayed rediploidization, all ohnologues are not ‘born equal’ and many will have opportunities to functionally diverge under unique environmental and ecological contexts, for example, during different phases of Earth’s climatic and biological evolution in the context of salmonid evolution [32]. It is also notable that ohnologues retained in LORe and null regions of the genome are enriched for different functions (Table S2), suggesting unique roles in adaptation, similar to past conclusions gained from comparison of ohnologues vs. small-scale gene duplicates (e.g. [58-59]). However, LORe is quite distinct from small-scale duplication, considering that large blocks of genes with common rediploidization histories will get the chance to diverge in functions in concert, meaning selection on duplicate divergence can operate on a multi-genic level.

LORe is possible whenever speciation precedes (or occurs in concert) to rediploidization (Fig. 1). This scenario is probable whenever rediploidization is delayed, most relevant for autotetraploidization events, which have occurred in plants [60], fungi [2] and unicellular eukaryotes [e.g. 25] and was the likely mechanism of WGD in the stem vertebrate and teleost lineages [35, 43, 61]. However, LORe is not predicted under a classic definition of allotetraploidization, as rediploidization is resolved immediately. Nonetheless, after some allotetraploidization events, the parental genomes have high regional similarity (i.e. segmental allotetraploidy [62]), allowing prolonged tetrasomic inheritance in some genomic regions, leading to potential for LORe. Interestingly, past studies have provided indirect support for LORe outside salmonids, including following WGD in the teleost ancestor [43]. A recent analysis of duplicated Hox genes from the lamprey *Lethenteron japonicum* failed to provide evidence of 1:1 orthology comparing jawed and jawless vertebrates, leading to the radical suggestion of independent, rather than ancestral vertebrate WGD events [63]. However, if rediploidization was delayed until after the divergence of these major vertebrate clades, which occurred no more than 60-100 Myr after the common vertebrate ancestor split from ‘unduplicated’ chordates [64-65], such findings are parsimoniously explained by LORe. In other words, WGD events may be shared by all vertebrates [61, 66], but some ohnologues became diploid independently in jawed and jawless lineages. Gaining unequivocal support for LORe beyond salmonids will require careful phylogenomic approaches akin to those employed here.

Our findings also reveal a possible mechanism to explain why some lineages experienced delayed post-WGD species radiations i.e. the WGD Radiation lag-time model [30-32]. This is a topical subject, given the recent finding that teleosts radiated at a similar rate to their sister lineage (holosteans) in the immediate wake of the teleost-specific WGD (Ts3R) [27], but nonetheless experienced much later radiations [28-29]. Our results suggest that in the presence of delayed rediploidization, the functional outcomes of WGD need not arise ‘explosively’, but can be mechanistically delayed for tens of Myr. For example, tissue expression responses for master genes required for saltwater tolerance are evidently fulfilled by one member of a salmonid ohnologue pair that began to diverge in functions 40-50 Myr post-WGD (Fig. 7). Hence, in light of evidence for delayed rediploidization after Ts3R [43], an alternative hypothesis is that teleosts gained an increasing competitive advantage through time compared to their unduplicated sister group, via the drawn-out creation of functionally-divergent ohnologue networks that provided greater scope for adaptation to ongoing environmental change. Similar arguments apply for delayed radiations in angiosperm lineages sharing WGD with a sister clade that diversified at a lower rate [30-31], offering a worthy area of future investigation.

For salmonids, climatic cooling likely provided a key selective pressure promoting the lineage-specific evolution of anadromy, which facilitated higher speciation rates in the long-term [32]. The elevated rediploidization rate in Salmoninae around the time that anadromy evolved, coupled to the lineage-specific regulatory divergence of LORe ohnologues regulating smoltification (Fig. 5, 7), allows us to hypothesize that LORe contributed to the evolution of lineage-specific adaptations that promoted species radiation. However, the role of LORe in adaptation is likely complex, occurring in a genomic context where an existing substrate of ‘older’ ohnologues (which by definition have had greater opportunity to diverge in functions) can also contribute to lineage-specific adaptation. This is evident in our data, as many relevant ohnologues from null regions of the genome show extensive regulatory divergence in the context of smoltification (Fig. 7; Dataset S1). A realistic scenario for lineage-specific adaptation involves functional interactions between networks of newly-diverging LORe ohnologues and older ohnologues that have already diverged from the ancestral state. Nonetheless, even though all ohnologues may undergo lineage-specific functional divergence, only during the initial stages of LORe will neofunctionalization and subfunctionalization [6-7, 11] arise without the influence of selection on past functional divergence (Fig. 1). In the future, follow up questions on the roles of LORe and null ohnologues (and their network-level interactions) in lineage-specific adaptation will become possible through comparative analysis of multiple salmonid genomes, done in a phylogenetic framework spanning the evolutionary transition to anadromy [67].

## Conclusions

LORe has unappreciated significance as a nested component of genomic and functional divergence following WGD and should be considered within investigations into the role of WGD as a driver of evolutionary adaptation and diversification, including delayed post-WGD radiations.

## Methods

### Target-enrichment and Illumina sequencing

To generate a genome-wide ohnologue set for phylogenomic analyses in salmonids, we used in-solution sequence capture with the Agilent SureSelect platform prior to sequencing on an Illumina HiSeq2000. Full methods were recently detailed elsewhere, including the source and selection of 16 study species [46]. While this past study was a small-scale investigation of a few genes [46], here we up-scaled the approach to 1,294 unique capture probes (Table S5; Text S3 provides details on probe design). 120mer oligomer baits were synthesised at fourfold tiling across the full probe set and a total of 1.5Mbp of unique sequence data was produced in each capture library. The captures were performed on randomly-fragmented gDNA libraries, meaning that the recovered data represent exons plus flanking genomic regions [46]. We recovered 21.7 million reads per species on average after filtering low-quality data (SD: 0.8 million reads; >99.1% paired-end data; see Table S6), which were assembled using SOAPdenovo2 [68] with a K-mer value of 91 and merging level of 3 (otherwise default parameters). Species-specific BLAST databases [69] were created for downstream analyses. Assembly statistics were assessed via the QUAST webserver [70] (Table S6). We used BLAST and mapping approaches to confirm that the sequence capture worked efficiently with high specificity and that pairs of ohnologues had been routinely recovered, even when a single ohnologue was used as a capture probe (see Text S3).

### Phylogenomic analyses

This work was split into a genome-wide investigation and a detailed study of Hox clusters. For both approaches, sequence data was sampled from our capture databases for different salmonid spp. using BLASTn [69] and aligned with MAFFT v.7 [71]. Northern pike was as used the outgroup to the Ss4R WGD in all analyses; this species was included in our target-enrichment study, but pike sequences were captured slightly less efficiently compared to salmonids [46]. Thus, we supplemented pike sequences using the latest genome assembly [47] (ASM72191v2; NCBI accession: CF_000721915). All phylogenetic tests were done at the nucleotide-level within the Bayesian Markov chain Monte Carlo (MCMC) framework BEAST v1.8 [72], specifying an uncorrelated lognormal relaxed molecular clock model [73] and the best-fitting substitution model (inferred by maximum likelihood in Mega v 6.0 [74] for individual alignments and PartitionFinder [75] for combinations of alignments). The MCMC chain was run for 10-million generations and sampled every 1,000th generation. TRACER v1.6 [76] was used to confirm adequate mixing and convergence of the MCMC chain (effective sample sizes >200 for all estimated parameters). Maximum clade credibility trees were generated in TreeAnnotator v1.8 [72]. All sequence alignments and Bayesian gene trees are provided in Table S1, including details on ohnologues sampled from the Atlantic salmon genome, alignment lengths and the best-fitting substitution model.

For the genome-wide study, the 1,294 unique capture probes were used in BLASTn searches against the Atlantic salmon genome (ICSASG_v2; NCBI accession: GCF_000233375) via http://salmobase.org/. This provided a genome-wide overview of the location of ohnologue alignments that could be generated via our capture assemblies and confidence that the targeted genes were true ohnologues retained from the Ss4R WGD, based on their location with collinear duplicated (homeologous) blocks [38]. In total, 384 ohnologue alignments were generated sampling regularly across all chromosomes, using the appropriate probes as BLAST queries against our capture databases to acquire data. Each tree contained a pair of verified ohnologues from Atlantic salmon and putative ohnologues captured from at least one species per each of the most distantly-related lineages within the three salmonid subfamilies.

For the Hox study, we used 89 Hox genes from Atlantic salmon [53] as BLASTn queries against our capture assemblies. The longest captured regions were aligned, leading to 54 alignments (accounted for within the 384 ohnologue alignments mentioned above) spanning all characterized Hox clusters [53]. We performed individual-level phylogenetic analyses on each dataset, revealing a highly congruent phylogenetic signal across different Hox genes from each Hox cluster (Fig. S2-S8), allowing alignments to be combined to the level of whole Hox clusters. To estimate the timing of rediploidization in the duplicated HoxBa cluster of salmonids [49], we employed the dataset combining all sequence alignments (i.e. tree in Fig. 4A). However, the analysis was done after setting calibration priors at four nodes according to MCMC posterior estimates of divergence times from a previous fossil-calibrated analysis [32]. The calibrations were made for the ancestor to two salmonid-specific HoxBa ohnologue clades for Salmoninae and Coregoninae. For Salmoninae, we set the prior for the common ancestor to *Hucho, Brachymystax, Salvelinus, Salmo* and *Oncorhynchus* (normally distributed, median = 32.5 Ma; SD: 3.5 Ma; 97.5% interval: 25-39 Ma). For Coregoninae, we set the prior for the common ancestor to *Stenodus leucichthys* and *Coregonus lavaretus* (normally distributed, median = 4.2 Ma; SD: 0.9 Ma; 97.5% interval: 2.4-5.7 Ma). We ran the calibrated BEAST analysis without data to confirm the intended priors were recaptured in the MCMC sampling.

Our estimates for the rate of ohnologue rediploidization in different salmonid lineages were made under the following assumptions: 1) that the common ancestor to salmonids had the same number of ohnologues as detected in regions of the Atlantic salmon genome defined by phylogenomic analysis (16,786 genes), which is a conservative estimate [38], 2) that 27% of these ohnologues (4,532 pairs) evolved under LORe (as defined for Atlantic salmon) and hence diverged independently in each salmonid subfamily, and 3) that 50%, 16% and 67% of LORe ohnologues (2,266, 725 and 3,036 pairs, respectively; estimated by gene tree topologies sampled across the genome, which should not be biased) experienced rediploidization during the time separating the ancestor of Salmoninae, Coregoninae and Thymallinae, from the crown of each subfamily, estimated at 19.5, 25.5 and 38.5 Myr, respectively [32].

### RNAseq analyses

To analyse ohnologue regulatory divergence in a relevant physiological condition to explore the evolution of anadromy, we performed RNAseq on 9 Atlantic salmon tissues sampled before and after smoltification. Six fish (three males and three females) were sampled from both freshwater (i.e. pre-smoltification, n=6; mean/SD length: 18.6/0.5cm) and saltwater (i.e. post-smoltification, n=6, mean/SD length: 25.8/0.8cm) at AquaGen facilities (Trondheim, Norway). RNA extraction was performed on each tissue and its purity and integrity was assessed using a Nanodrop 1000 spectrophotometer (Thermo-Scientific) and 2100 BioAnalyzer (Agilent), respectively. Subsequently, stranded libraries were produced from 2μg of total RNA using a TruSeq stranded total RNA sample Kit (Illumina, USA) according to the manufacturer’s instructions (Illumina #15031048 Rev.E). Sequencing was performed on a MiSeq instrument using a v3. MiSeq Reagent Kit (Illumina) generating 2x300 bp of strand-specific, paired-end reads. For each tissue, the sequenced individuals were pooled into 2 sets of 3 individuals of each sex in both freshwater and saltwater (hence, any reported responses are common to males and females). For the global analysis of ohnologue expression divergence in different tissues under controlled conditions (i.e. Fig. 6A, B), we employed Illumina transcriptome reads previously generated for 15 Atlantic salmon tissues [described in 38].

In both RNAseq analyses, raw Illumina reads were subjected to adapter and quality-trimming using cutadapt [77], followed by quality control with FastQC, before mapping to the RefSeq genome assembly (ICSASG_v2) using STAR v2.3 [78]. Uniquely-mapped reads were counted using the HTSeq python script [79] in combination with a modified RefSeq .gff file. The .gff file was modified to contain the attribute “gene_id” (file accessible at http://salmonbase.org/Downloads/Salmo_salar-annotation.gff3). Expression levels were calculated as counts per million total library counts in EdgeR [80]. Total library sizes were normalised to account for bias in sample composition using the trimmed mean of m-values approach [78]. For the smoltification study, log-fold expression changes were calculated, contrasting samples from freshwater and saltwater, done separately for each tissue using EdgeR [80]. Genes showing a FDR-corrected *P* value ≤ 0.05 were considered differentially expressed.

To identify salmonid-specific ohnologue pairs in null and LORe regions of the Atlantic salmon genome, a self-BLASTp analysis was done using all annotated RefSeq proteins - keeping only proteins coded by genes within verified collinear (homeologous) regions retained from the Ss4R WGD [38] with >50% coverage and > 80% identity to both query and hit. Statistical analyses on expression data were performed using various functions within R [81]. Expression divergence was estimated using Pearson correlation in all cases. The Circos plot (Fig. 6A) was generated using the circlize library in R [82].

### GO enrichment analyses

GO annotations for Atlantic salmon protein-coding sequences were obtained using Blast2GO [83]. The longest predicted protein for each gene was blasted against Swiss-Prot (http://www.ebi.ac.uk/uniprot) and processed with default Blast2GO settings [84]. The results have been bundled into an R-package (https://gitlab.com/Cigene/Ssa.RefSeq.db). Protein-coding genes were tested for enrichment of GO terms belonging to the sub-ontology ‘Biological Process’ using a Fisher test implemented in the Bioconductor package topGO [84]. The analysis was restricted to terms of a level higher than four, with more than 10 but less than 1,000 assigned genes. Enrichment analyses were done separately for all ohnologue pairs with annotations retained in LORe (2,002 pairs) and null (5,773 pairs) regions of the RefSeq genome assembly. We recorded the chromosomal locations of LORe ohnologues for the most significantly-enriched GO terms, including the number of unique LORe regions they occupy in the genome (Table S4). The rationale was to establish the extent to which ohnologues underlying an enriched GO term are physically clustered. We devised a ‘clustering index’, quantifying the total number of cases where n ≥ 2 ohnologues present within the relevant genomic regions are located ≤500Kb, expressed as a proportion of n-1 the total number of ohnologues located within those regions. A respective clustering index of 1.0, 0.5 and 0.0 means that all, half or zero of the retained ohnologues (accounting for an enriched GO term) are located within 500Kb of their next nearest gene within the same genomic region. 500Kb was considered a conservative distance to capture genes expanded by tandem duplication.

## Declarations

### Ethics approval and consent to participate

Atlantic salmon tissues were sampled by AquaGen (Trondheim, Norway) following Norwegian national guidelines with regard to the ethical aspects of animal welfare.

### Consent for publication

Not applicable

### Availability of data and material

Illumina sequence reads for the sequence capture study were deposited in NCBI (Bioproject: PRJNA325617). All sequence alignments and phylogenetic trees used for phylogenomics are provided in Table S1. Illumina sequence reads for the tissue expression study performed under controlled conditions can be found in the NCBI SRA database (accessions: SRX608594; SRS64003; SRS640030; SRS640015; SRS640003; SRS639997; SRS639041; SRS640021; SRS639990l; SRS639861; SRS640009; SRS639992; SRS639037; SRS640002; SRS639994). Illumina sequence reads for the fresh to saltwater transition experiment were deposited in the European Nucleotide Archive (project accession number: SRP095919).

### Author contributions

DJM and PWH defined the LORe model. DJM designed the sequence capture study. FML performed lab work for the sequence capture study and contributed to probe design. FML, MKG and DJM performed phylogenetic analyses. FG, TRH and SRS performed the expression analyses. FG performed GO enrichment analyses. DJM and MKG interpreted GO enrichment analyses. FML, MKG, FG, SRS and DJM designed figures/tables. DJM drafted the manuscript. All authors interpreted data and contributed to the writing of the final manuscript.

## Acknowledgements

We are grateful to Dr Steven Weiss (University of Graz, Austria), Dr Takashi Yada (National Research Institute of Fisheries Science, Japan), Dr Robert Devlin (Fisheries and Oceans Canada, Canada), Mr Neil Lincoln (Environment Agency, UK), Dr Kevin Parsons (University of Glasgow, UK), Prof. Colin Adams (University of Glasgow, UK) and Mr Stuart Wilson (University of Glasgow, UK) for providing salmonid material used in the sequence capture study or assisting with its collection. We thank staff at the Centre for Genomic Research (University of Liverpool, UK) for performing sequence capture and Illumina sequencing, Maren Mommens (AquaGen, Norway) for sampling salmon tissues and Hanne Hellerud Hansen (CIGENE, Norway) for performing laboratory work for RNAseq.

